# Nitrogen metabolism in the picoalga *Pelagomonas calceolata*: disentangling cyanate lyase function under different nutrient conditions

**DOI:** 10.1101/2024.02.19.580968

**Authors:** Nina Guérin, Chloé Seyman, Céline Orvain, Laurie Bertrand, Priscillia Gourvil, Ian Probert, Benoit Vacherie, Élodie Brun, Ghislaine Magdelenat, Karine Labadie, Patrick Wincker, Adrien Thurotte, Quentin Carradec

## Abstract

Cyanate (OCN^-^) is potentially an important organic nitrogen source in aquatic environments given the prevalence and activity of cyanate lyase genes in microalgae. However, the conditions under which these genes are expressed and the actual capacity of microalgae to assimilate cyanate remain underexplored. Here, we studied the nitrogen metabolism of the cosmopolitan picoalga *Pelagomonas calceolata* (Pelagophyceae, Stramenopiles) in environmental metatranscriptomes and transcriptomes from culture experiments under different nitrogen sources and concentrations. We observed that cyanate lyase is over-expressed in nitrate-poor oceanic regions, suggesting that cyanate is an important molecule contributing to the persistence of *P. calceolata* in oligotrophic environments. In the laboratory, we confirmed that this gene is over-expressed in low-nitrate medium together with several genes involved in nitrate recycling from endogenous molecules. Non-axenic cultures of *P. calceolata* were capable of growing on various nitrogen sources, including nitrate, urea and cyanate, but not ammonium. RNA sequencing of these cultures revealed that cyanate lyase was under-expressed in the presence of cyanate, indicating that this gene in not involved in the catabolism of extracellular cyanate to ammonia. Conversely, axenic *P. calceolata* cultures were not able to grow on cyanate, suggesting that the bacterial community consumes cyanate and provides an available form of nitrogen for growth of the alga. Based on environmental datasets and laboratory experiments, we propose that cyanate lyase is important in nitrate-poor environments to reduce the toxicity of intracellular cyanate produced by endogenous nitrogenous compound recycling, rather than being used to metabolise imported extracellular cyanate as an alternative nitrogen source.

## Introduction

Nitrogen is essential to many biological processes, such as photosynthesis, amino acid and nucleic acid biosynthesis, thus its bioavailability in the oceans impacts the growth of primary producers (Moore et al., 2013). The primary sources of fixed nitrogen for phytoplankton are inorganic ammonium (NH_4_ ^+^), nitrite (NO_2_ ^-^) and nitrate (NO_3_ ^-^) (Hutchins & Capone, 2022). Nitrate is the most abundant nitrogenous compound, whereas nitrite and ammonium are typically less abundant (Collos & Berges, 2002). Nevertheless, ammonium is considered a preferred source of nitrogen for phytoplankton due to its lower energy cost during assimilation (Kumar & Bera, 2020). The majority of oceanic surface waters are depleted in inorganic nitrogen compounds because of uptake by photosynthetic organisms in the photic zone (Hutchins & Capone, 2022). One consequence of ocean warming is enhanced stratification, which reduces the supply of nutrients to the euphotic zone (Fu et al., 2016). By the end of the twenty-first century, an average 1.06 ± 0.45 mmol.m^-3^ decrease in nitrate concentration in surface waters is projected under the IPCC high-emission scenario SSP5-8.5 (Kwiatkowski et al., 2020).

Since inorganic nitrogen compound concentrations are variable, photosynthetic organisms have developed a range of strategies to face temporal and spatial fluctuations in nitrate, nitrite and ammonium availability. Optimisation of inorganic nitrogen uptake can be achieved by regulation of the expression of transporters in several phytoplankton groups, notably prasinophytes and diatoms (Lampe et al., 2019). Storage and recycling of nitrogen-rich proteins is also an important strategy in diatoms (Alipanah et al., 2015; Scarsini et al., 2022). Various phytoplankton can metabolise dissolved organic nitrogen (DON) compounds such as urea, purines or amino acids (Kumar & Bera, 2020). In the ocean, the concentration of DON, often higher than that of dissolved inorganic nitrogen (DIN), contributes to autotrophic production and growth of primary producers, especially in coastal and estuarine environments (Sipler & Bronk, 2015). DON compounds have been shown to support the growth of microalgae in low N-conditions. For example, urea can be used as a nitrogen source by many phytoplankton groups, such as diatoms, dinoflagellates and the bloom-forming pelagophyte *Aureococcus anophagefferens* (Berg et al., 1997; Cerón GarcÍá et al., 2005; García-Portela et al., 2020). Diatoms such as *Phaeodactylum tricornutum* and *Thalassiosira pseudonana* possess urea transporters that are overexpressed in N-limited conditions. The coccolithophore *Emiliania huxleyi* can use a broad range of organic nitrogen sources, including urea, hydroxyurea, hydroxanthine, purines and small amides such as acetamides and formamides (Palenik & Henson, 1997).

The cyanate ion (OCN^-^), which is the smallest nitrogenous organic compound, was originally described as a toxic molecule altering structural and functional properties of proteins through carbamylation (Jaisson et al., 2011). In the oceans, cyanate originates from terrestrial inputs, spontaneous decomposition of carbamoyl-phosphate or urea released by zooplankton or senescent phytoplankton, as well as via photochemical degradation of dissolved organic matter (Wang et al., 2024; Widner et al., 2016). Cyanate concentrations up to 45 nM have been recorded in subsurface ocean waters (Widner et al., 2018). The organisms capable of cyanate uptake and metabolism in the environment, as well as the underlying molecular mechanisms for this process, remain unclear. Growth on cyanate as the sole nitrogen source was first described in *Escherichia coli*, then in cyanobacteria, the ammonia-oxidizing archaea *Nitrososphaera gargensis*, several yeasts, and the mixotrophic dinoflagellate *Prorocentrum* (Anderson et al., 1990; Hu et al., 2012, 2014; Kamennaya & Post, 2011; Linder, 2018; Palatinszky et al., 2015). In bacteria, the cyanate lyase enzyme (cynS gene) catalyses the bicarbonate-dependant break down of cyanate to ammonia and carbon dioxide (Johnson & Anderson, 1987). Cyanate lyase homologs are present in the genome of many organisms, including dominant eukaryotic phytoplankton lineages (Mao et al., 2022). In the environment, the expression of cyanate lyase in most prokaryotic and eukaryotic phytoplankton is increased in N-limited environments, suggesting that these organisms might use cyanate as an alternative nitrogen source (Dong et al., 2014; Mao et al., 2022; Smith et al., 2019; Wurch et al., 2011). ^15^N stable isotope probing revealed that cyanate uptake could account for up to 10% of total nitrogen uptake in natural communities from the offshore oligotrophic Atlantic, particularly in surface waters (Widner et al., 2016). Since cyanate lyase expression is as important as urease expression in the environment, it has been suggested that cyanate has a crucial ecological role (Mao et al., 2022). However, the uptake of cyanate in eukaryotic phytoplankton has only been demonstrated in *Prorocentrum*. A recent study reported that cyanate enrichment in natural phytoplankton populations induces growth of the picocyanobacterium *Synechococcus*, but not that of eukaryotic phytoplankton (Sato et al., 2023). This leads us to question the role of cyanate lyase in organic nitrogen assimilation in photosynthetic eukaryotes.

In order to disentangle the endogenous role of cyanate lyase in microeukaryotes and more generally to develop a better understanding of eukaryotic phytoplankton acclimation to varying nitrogen availability in the environment, we used the cosmopolitan photosynthetic picoeukaryote *Pelagomonas calceolata* (Pelagophyceae) as a model for pelagic phytoplankton (Guérin et al., 2022; Worden et al., 2012). The *P. calceolata* (strain RCC100) genome contains a large set of genes involved in nitrogen metabolism (Guérin et al., 2022). The presence of genes coding for arginase, urease and cyanate lyase may suggest the capacity for metabolism of organic nitrogen compounds. In the environment, *P. calceolata* overexpresses nitrogen ion transporters, nitrate and nitrite reductases, glutamine synthetases, nitrate/nitrite sensing proteins and cyanate lyase in low-N environments (Dupont et al., 2015; Guérin et al., 2022). Here we cultivated two *P. calceolata* strains under high or limited nitrate conditions and tested the effect of several inorganic (nitrate, ammonia) and organic (urea, cyanate) sources of nitrogen on *P. calceolata* growth. For all of these conditions, we performed RNA sequencing and identified differentially expressed genes. We compared our results with a differential analysis of *P. calceolata* gene expression levels in environmental metatranscriptomes from the *Tara* Oceans expedition according to *in situ* measurements of nitrate concentrations.

## Materials & Methods

### Environmental metatranscriptomes and associated metadata

Metatranscriptomic datasets from *Tara* Oceans and *Tara* Polar Circle expeditions (Alberti et al., 2017) were used to detect *in situ* gene expression of *P. calceolata*. All available datasets from seawater samples of the photic zone (73 surface and 51 deep-chlorophyll maximum samples) and from 2 size-fractions (80 from the 0.8-5 µm fraction and 44 from the 0.8-2000 µm fraction) were selected. Metatranscriptomic reads were aligned on the 16,667 predicted mRNA sequences of the *P. calceolata* genome (ENA, PRJEB47931) with bwa-mem2 version 2.2.1 with default parameters (Li et al., 2009). Reads aligned on the *P. calceolata* genome with >95% identity over 80% of read length were selected. Nuclear genes covered by a minimum of 10 reads in at least 10 samples were retained. To eliminate putative cross-mapped genes (i.e. highly conserved genes which probably aggregate reads from other organisms), genes detected in >90% of samples (including those where *P. calceolata* is not present) were removed. Finally, only samples with >75% of *P. calceolata* genes were kept for the next steps. The environmental parameters measured during the expedition are available in the Pangaea database (https://www.pangaea.de/) (Pesant et al., 2015). Nitrate concentrations were calculated from *in situ* sensor (SATLANTIC) data, calibrated using water samples. Samples were considered “low-nitrate” if they contained <2 μM of nitrate. Differential expression analyses were conducted with DESeq2 package version 1.32.0 under R version 4.1.1 across the 15,617 genes and the 112 environmental samples with available *in situ* nitrate concentration measurements (Love et al., 2014). Pairwise comparisons were performed across 43 “high-nitrate” samples (nitrate concentration > 2 µM) and 69 “low-nitrate” samples (nitrate concentration < 2µM) and with the function DESeq with default parameters and log2 fold change values were calculated with lfcShrink function. Genes with a p-value < 0.01 and a log2 fold change > 2 or < -2 were considered as differentially expressed.

### *P. calceolata* cultures in different nitrogen conditions

*Pelagomonas calceolata* strains RCC100 and RCC697 (obtained from the Roscoff Culture Collection: www.roscoff-culture-collection.org) were cultivated in artificial seawater (ASW) supplemented by L1 medium (as described in Guillard and Hargraves 1993). ASW was prepared by dissolution of 24.55g of sodium chloride (NaCl), 0.75g of potassium chloride (KCl), 4.07g of magnesium chloride hexahydrate (MgCl_2_*6 H2O), 1.11g of calcium chloride (CaCl_2_), 2.95g of magnesium sulphate (MgSO_4_) and 0.21g of sodium bicarbonate (NaHCO_3_) in 1L of sterile distilled water. 1 ml of trace metals, vitamins and nutrients from the Bigelow L1 medium Kit (MKL150L) were added to attain the following concentrations: 882 µM sodium nitrate (NaNO_3_), 36.2 µM monosodium phosphate (NaH_2_PO_4_ ^-^) and 106 µM sodium silicate (Na_2_SiO_3_). Cultures were maintained at 20°C under a 12:12h light-dark photoperiod and a blue light (while LEDs covered by a blue filter: LEE FILTERS, 183 Moonlight blue) at an intensity of 20 μmol. m^−2^.s^−1^ of photosynthetic photons. The non-flagellated *P. calceolata* strain (RCC697) was maintained on an orbital shaker (Kühner) at 150 r.min^-1^. For the nitrate depletion experiment, the nitrate concentration of the L1 medium was reduced to 441 µM, 220 µM, 110 µM or 50 µM. For the cyanate and the ammonium experiments, nitrate was replaced by potassium cyanate (KOCN) or ammonium chloride (NH_4_Cl) at the same concentration (882 µM). For the urea experiment, nitrate was replaced by urea (CH4N2O) at 441 µM to keep the same concentration of nitrogen atoms in all conditions.

To obtain axenic *P. calceolata* cells (Figure 3b), RCC100 was preliminarily treated with a mix of antibiotics (Spectinomycin (50 µg/ml), Neomycin (100 µg/ml) and Carbenicillin (30 µg/ml). Axenicity was verified on marine broth plates (Difco 2216). No reduction in growth of axenic vs non-axenic cultures was observed under standard culture conditions. During this experiment, *P. calceolata* cells were counted daily with a flow cytometer (Cytoflex, Beckman Coulter Life Sciences). Cell counts were fitted to a logistic growth curve with the R package Growthcurver (version 0.3.1). The r statistics given by GrowthCurver correspond to the growth rate that would occur if there were no restrictions imposed on total population size.

**Figure 1.**
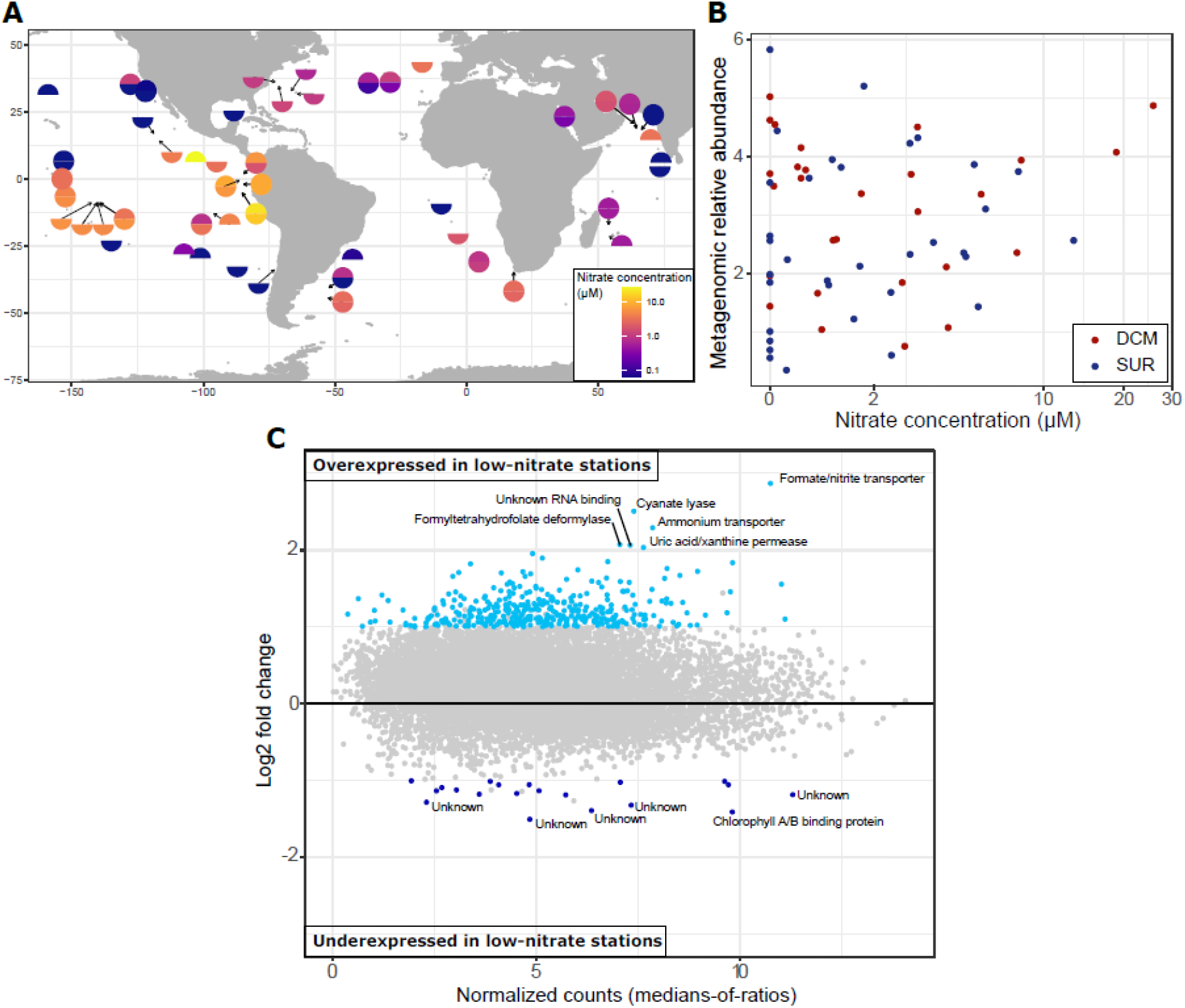
Abundance and transcriptomic response of *P. calceolata* to environmental nitrate concentrations. A) Nitrate concentrations measured during the *Tara* Oceans expedition. The colour code indicates nitrate concentrations in µmol/l for surface and DCM samples in the upper and lower part of each dot respectively. B) Relative abundance of *P. calceolata* in *Tara* samples estimated from metagenomics reads according to the concentration of nitrate (µM) C) *P. calceolata* gene expression levels between low-nitrate (NO_3_^-^ < 2 µM, n=69) and high-nitrate samples (NO_3_^-^ > 2 µM, n=43). Log2FC between low- and high-nitrate samples are given according to their mean expression level (normalized with DESeq2). Differentially expressed genes with p-value < 0.01 and log2FC >1 or < -1 are coloured in blue.

**Figure 2:**
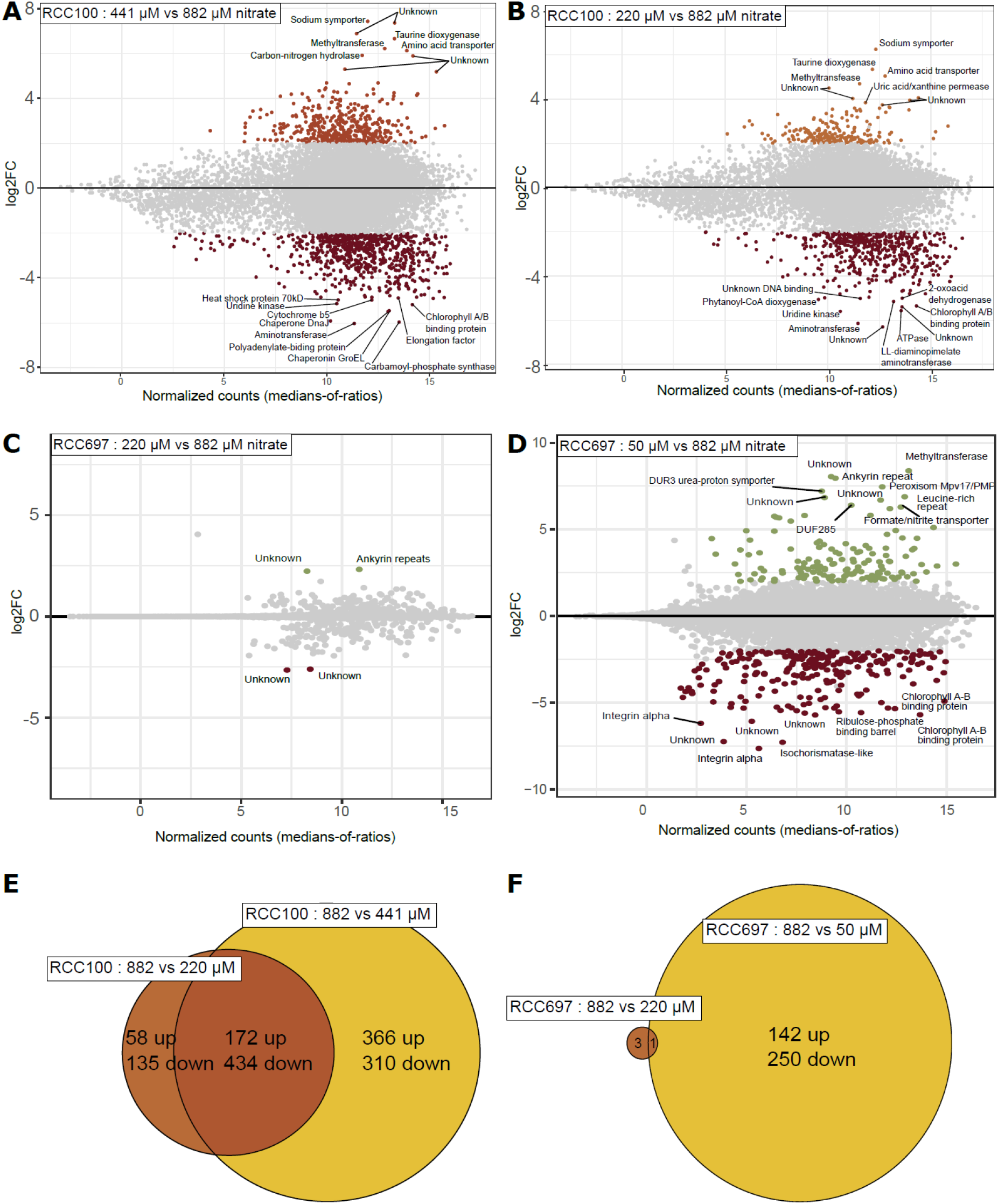
Transcriptomic response of *P. calceolata* to low-nitrate culture conditions. A-D) Differentially expressed genes of *P. calceolata* RCC100 in 441 µM (A) and 220 µM (B) nitrate and RCC697 in 220 µM (C) and 50 µM (D) compared to 880 µM nitrate. Genes with p-value < 0.01 and log2FC > 2 are coloured. D, E) Euler diagram of DEGs in RCC100 (E) and RCC697 (F). The number of upregulated and downregulated genes is indicated.

**Figure 3:**
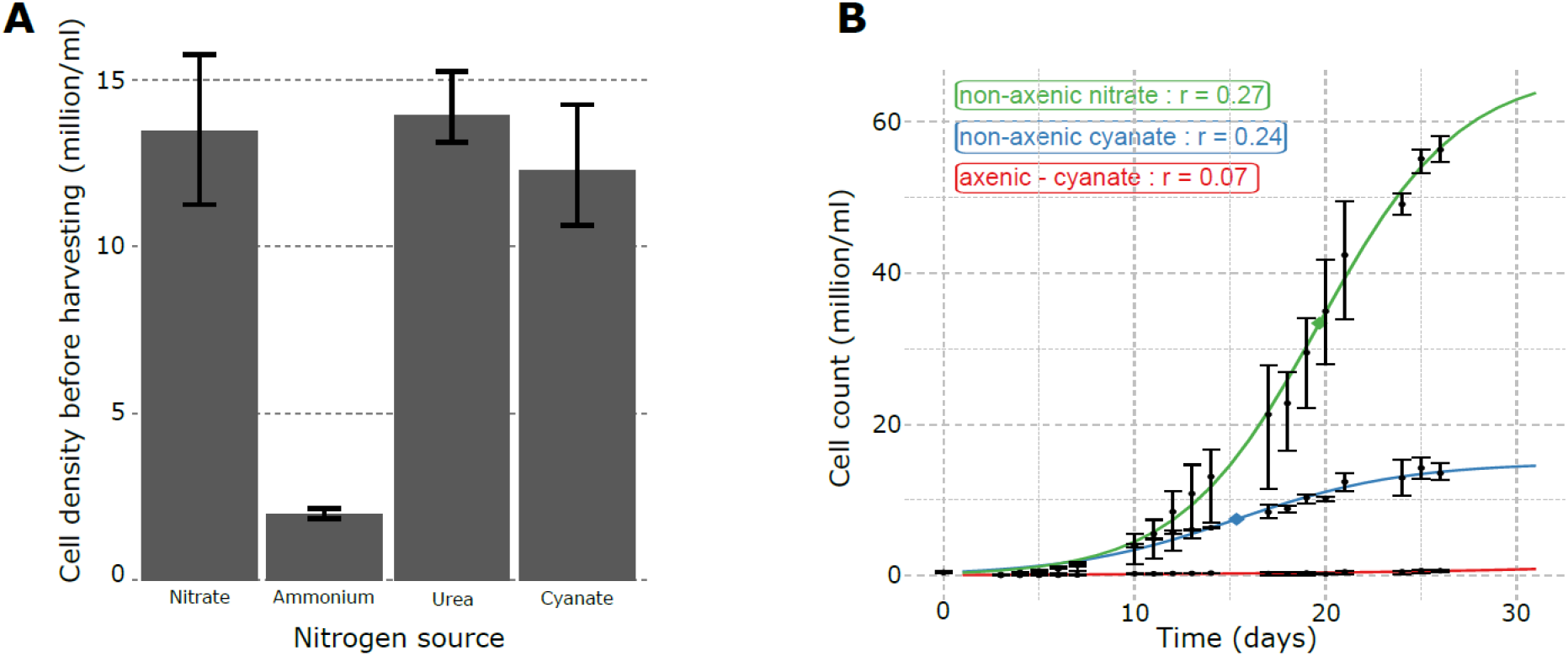
*P. calceolata* growth with different nitrogen sources. A) Average *P. calceolata* density (cell/ml) before harvesting for RNA extraction. Cells counts were measured on a Thoma counting chamber under light microscopy. B) *P. calceolata* growth cultivated with nitrate, cyanate or cyanate after antibiotic treatments to remove the bacterial community. Cell counts were estimated with a flow cytometer. Experiments in A and B were carried out in triplicate and error bars indicate minimum and maximum values for each condition.

RCC100 and RCC967 were grown in each condition tested during an acclimation phase lasting a minimum of 8 days. Cultures were then diluted in fresh medium in triplicate and grown for 5 to 9 days until they reached a minimum concentration of 10 million cells per ml. Growth and fluorescence were monitored daily using a Qubit instrument (Invitrogen™ Qubit™ 3 Fluorometer Q33216, blue excitation at 470 nm, far red emission 665-720 nm) and cells were counted on a Thoma cell (Marienfeld Thoma counting chamber, depth 0.1 mm, 0640710) under a microscope on the day of harvesting. *P. calceolata* cells were harvested by filtration through 1.2 µm mixed cellulose-ester membrane filters (MF-Millipore, rawp04700) with a peristaltic pump (SFP-100), then transferred into 15ml tubes, flash-frozen in liquid nitrogen, and stored at -80°C until RNA extraction.

### RNA extraction and sequencing

Flash-frozen filters were vortexed in QIAzol, then RNA was extracted using RNeasy Plus Universal Mini Kits (Qiagen, Ref 73404) following the manufacturer’s instructions. All extracted RNA samples were treated with 6U of TURBO™ DNase (2 U/µl) (Thermo Fisher Scientific, Ref. AM2238) then purified with RNA Clean and Concentrator-5 kit (Zymo Research, Ref. ZR1016), keeping only large RNA fractions (> 200 nt) for RNAseq library preparation. 100 ng of treated RNA were used to produce Illumina libraries (Illumina Stranded mRNA Prep, Ligation). Briefly, poly(A) + RNAs were selected with oligo(dT) beads, chemically fragmented by divalent cations under high temperature, converted into single-stranded cDNA using random hexamer priming, followed by second strand synthesis and 3′-adenylation. A pre-index anchor was ligated and a PCR amplification step with 15 cycles was conducted to add 10bp unique dual index adapter sequences (IDT® for Illumina® RNA UD Indexes, Ligation). All libraries were quantified by Qubit dsDNA HS Assay measurement. A size profile analysis was performed in an Agilent 2100 Bioanalyzer (Agilent Technologies, Santa Clara, CA, USA). The library preparation failed for one sample of RCC697 (200 µM NO_3_). Libraries were sequenced in 2×150 bp on an Illumina NovaSeq 6000 sequencer (Illumina, San Diego, CA, USA) in order to obtain 50 million paired-end reads. After Illumina sequencing, an in-house quality control process was applied to the reads that passed the Illumina quality filters (Alberti et al., 2017). Briefly, Illumina sequencing adaptors and primer sequences were removed, then low-quality nucleotides (Q < 20) were discarded from both ends of the reads. Sequences between the second unknown nucleotide (N) and the end of the read were also trimmed. Reads shorter than 30 nucleotides were discarded after trimming with an adaptation of the fastx_clean tool (https://www.genoscope.cns.fr/fastxtend/). In the last step, reads that were mapped to the Enterobacteria phage PhiX174 genome (GenBank: NC_001422.1) were discarded using bowtie2 v2.2.9 (-L 31–mp 4–rdg 6,6–local–no-unal) (Langmead & Salzberg, 2012). Remaining rRNA reads were removed using SortMeRNA v2.1 and SILVA databases (Kopylova et al., 2012; Quast et al., 2013).

### Analysis of gene expression levels in different nitrogen conditions

As for analysis of environmental metatranscriptomes, RNAseq reads were aligned with bwa-mem2 version 2.2.1 on the *P. calceolata* genome. Reads with a minimal size of 50 bp and aligned with >95% identity over 80% of read length were selected. 16,659 out of 16,667 genes were detected in at least one sample. Only nuclear genes were retained, with gene expression levels normalised in transcripts per kilobase per million mapped reads (TPM). Pearson’s correlation matrices were computed with R using cor function between triplicates and across conditions, based on the gene expression normalised by TPM. Hierarchical clustering of Euclidian distance between samples was performed with dist and hclust functions of stats package on R version 4.1.1. Differential expression analysis on transcriptomic samples was carried out in the same way as for the metatranscriptomic DESeq2 analysis above. Identification of differentially expressed genes between control and test samples was performed by pairwise comparisons across the standard condition (882 µM NO3) and low-nitrate conditions (220 µM or 441 µM NO3) on the one hand, and across the standard condition and changing nitrogen sources (882 µM ammonium, 882 µM cyanate and 441 µM urea) on the other hand. Genes presenting a p-value < 0.01 and a log2 fold change > 2 or < -2 were considered as differentially expressed. We specifically looked for log2 fold change of genes identified as involved in the nitrogen cycle in our previous study (Guérin et al., 2022). Figures 1 to 5 were generated with ggplot2 version 3.5.0 except for the Euler diagrams that were made with eulerr version 7.0.2 and graphics version 4.1.1.

### Functional re-annotation of *P. calceolata* genes

Functional annotation *P. calceolata* genes was updated for this study using InterProScan v5.61.93.0 (Quevillon et al., 2005) and the following protein domain databases : Pfam, Gene3D, TIGRfam, SMART, CDD and Gene Onthology. All matches with a pvalue below 1×10^-5^ were retained. A protein alignment against the NR database (24-08-2023 version) was performed with diamond v2.1.2 (Buchfink et al., 2021). The best match was retained if the e-value was below 1×10^-5^. The HMM search tool KofamKoala v1.3.0 was used to identify KEGG Orthologues (version of November 2023) (Aramaki et al., 2020). Annotations with an e-value < 1×10^−5^ and a score above the HMM threshold were retained. Homologies with protein clusters of the Eggnog database were identified with the eggnog-mapper tool version 2.1.12 using the *very-sensitive* mode and diamond aligner (Cantalapiedra et al., 2021; Huerta-Cepas et al., 2019). For the prediction of protein localization, DeepLoc version 2.0 (Thumuluri et al., 2022) and TargetP version 2 in eukaryote mode (Almagro Armenteros et al., 2019) were used. This methodology was applied on the 16,667 gene models, translated in the 6 frames in full with the transeq function of Emboss version 6.6 and on the protein predicted by Gmove (Dubarry et al., 2016). If functional annotations were identified on several frames, the frame with the best score was retained. This new version of the functional annotation of *P. calceolata* genes is available on github https://github.com/institut-de-genomique/PelagomonasNitrogenMetabolism/.

## Results

### In situ gene expression levels of P. calceolata according to nitrate concentration

To determine the *in situ* response of *P. calceolata* to low nitrogen concentration, we used all metatranscriptomes of *Tara* Oceans datasets (Alberti et al., 2017). We aligned metatranscriptomic reads on the predicted mRNAs of the *P. calceolata* RCC100 genome and selected 124 *Tara* samples where at least 75% of the genes were detected (at least 1 read aligned with more than 95% identity over 80% of its length). For 112 of these samples with available *in situ* measurements of nitrate, 69 have nitrate concentrations below 2 µM and are considered “low-nitrate”, and 43 have nitrate concentrations above 2 µM and are considered “high-nitrate” (Figure 1A, Supplementary Table 1). We observed no significant correlation between nitrate concentrations and the relative abundance of *P. calceolata* (Figure 1B ; person= 0.2, p.value=0.11).

**Table 1:**
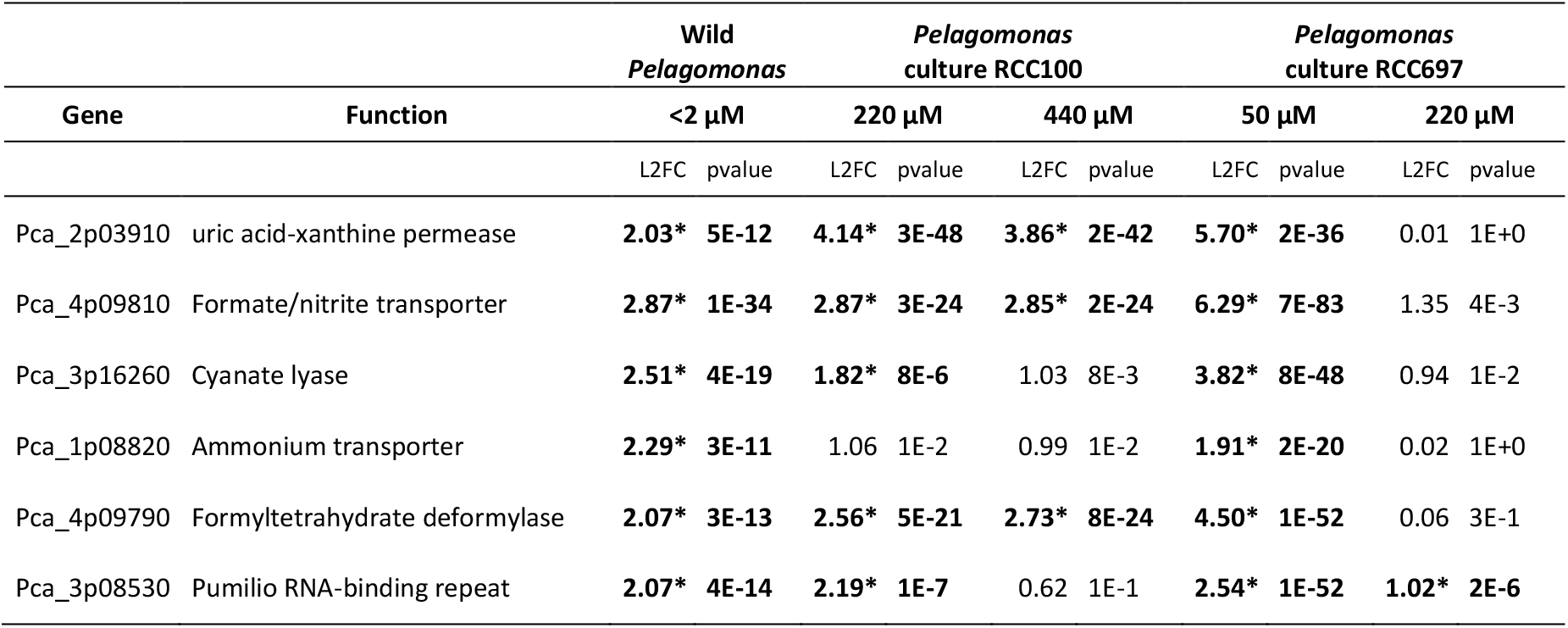
Differentially expressed genes in low nitrate conditions in the environment and in culture. Log2 fold changes for each condition and each gene overexpressed in low-nitrate environments are indicated. The complete list of DEGs in culture experiments is in Supplementary Table 4. *= indicates pvalues < 0.001.

| Gene | Function | Wild<br>*Pelagomonas* |  | *Pelagomonas*<br>culture RCC100 |  |  |  | *Pelagomonas*<br>culture RCC697 |  |  |  |
| --- | --- | --- | --- | --- | --- | --- | --- | --- | --- | --- | --- |
| | | <2 μM | | 220 μM | | 440 μM | | 50 μM | | 220 μM | |
|  |  | L2FC | pvalue | L2FC | pvalue | L2FC | pvalue | L2FC | pvalue | L2FC | pvalue |
| Pca_2p03910 | uric acid-xanthine permease | 2.03* | 5E-12 | 4.14* | 3E-48 | 3.86* | 2E-42 | 5.70* | 2E-36 | 0.01 | 1E+0 |
| Pca_4p09810 | Formate/nitrite transporter | 2.87* | 1E-34 | 2.87* | 3E-24 | 2.85* | 2E-24 | 6.29* | 7E-83 | 1.35 | 4E-3 |
| Pca_3p16260 | Cyanate lyase | 2.51* | 4E-19 | 1.82* | 8E-6 | 1.03 | 8E-3 | 3.82* | 8E-48 | 0.94 | 1E-2 |
| Pca_1p08820 | Ammonium transporter | 2.29* | 3E-11 | 1.06 | 1E-2 | 0.99 | 1E-2 | 1.91* | 2E-20 | 0.02 | 1E+0 |
| Pca_4p09790 | Formyltetrahydrate deformylase | 2.07* | 3E-13 | 2.56* | 5E-21 | 2.73* | 8E-24 | 4.50* | 1E-52 | 0.06 | 3E-1 |
| Pca_3p08530 | Pumilio RNA-binding repeat | 2.07* | 4E-14 | 2.19* | 1E-7 | 0.62 | 1E-1 | 2.54* | 1E-52 | 1.02* | 2E-6 |

Differential expression analysis between “high-nitrate” and “low-nitrate” environments revealed 375 genes significantly overexpressed in low-nitrate samples (p-value < 0.01), 6 of which had a log2 Fold Change (log2FC) higher than 2 (Figure 1B and Supplementary Table 2). Two of these genes are involved in inorganic nitrogen transport across cell membranes, a formate or nitrite transporter (PF01226 domain) and an ammonium transporter (PF00909 domain) as previously described (Guérin et al., 2022). In addition, a purine transporter (uric acid/xanthine permease, K23887 domain), a formyltetrahydrofolate deformylase (10-FDF; K01433-EC 3.5.1.10), an enzyme acting on carbon-nitrogen bonds and potentially related to ammonium recycling from glycine, as well as an unknown gene carrying a RNA-binding domain of the Pumilio family (K17943 domain) were overexpressed in low-nitrate environments. The cyanate lyase (PF02560 domain) was the second most overexpressed gene in the *P. calceolata* genome in low-nitrate environments. We also noted the slight overexpression of a nitrate/nitrite transporter (K02575 domain), a carbon nitrogen hydrolase (PF00795 domain) and a dipeptidase (K08659 domain), suggesting active recycling of nitrogen-rich molecules. Only 20 genes were slightly underexpressed in low-nitrate environments with a log2FC between 1 and 1.5, most of which (14 genes) have unknown functions. Three Light Harvesting Complex proteins (LHC; PF00504 domain) were downregulated in low-nitrate environments, suggesting reduced chloroplast activity.

### Gene expression variations under low-N conditions

To estimate acclimation capacities of *P. calceolata* to low-nitrate conditions, we cultivated two strains of this species complex. RCC100 (=CCMP1214), isolated in the Pacific Ocean in 1973, is flagellated with a cell size of 1.5-2 µm, while RCC697, isolated in the Indian Ocean in 2003, is significantly larger with a cell diameter of 3 µm and is non-flagellated (Supplementary Fig. 1 A and B). Since their isolation, both strains have been maintained in the Roscoff Culture Collection at 20°C. These strains were cultivated with different nitrate concentrations ranging from 50 µM to 882 µM for a minimum of 3 weeks with weekly dilutions in fresh media. The minimal nitrate concentration to observe cell growth and reach a sufficient number of cells for RNA extraction was 220 µM for RCC100 and 50 µM for RCC697.

*P. calceolata* cells were harvested in exponential growth phase for RNA extraction in the following conditions: 220, 441 and 882 µM for RCC100 and 50, 220, and 880 µM for RCC697. Between 29.9 and 84.5 million paired-end RNA reads were sequenced for each sample, then aligned on the predicted mRNAs of the *P. calceolata* RCC100 genome. 86% of RCC100 genes were covered by at least one RCC697 read with on average 96.0% identity, demonstrating the genetic proximity of the two strains (Supplementary Table 3). Correlations of gene expression levels within triplicates were higher (Pearson’s r coefficients > 0.95) than correlations between conditions (Pearson < 0.81), with the exception of RCC697 882 µM vs RCC697 220 µM nitrate, that were highly correlated (Pearson > 0.92). These correlations indicate that all tested conditions induced a transcriptomic response of *P. calceolata*, except the reduction to 220 µM nitrate in RCC697 (Supplementary Figure 2).

Differential expression analyses were computed between the high-nitrate condition (882 µM) and each tested condition for both strains. For RCC100, 1,282 genes were differentially expressed with 441 µM nitrate and 799 were differentially expressed with 220 µM nitrate (Figure 2 A and B). A large proportion of these differentially expressed genes (DEGs) were common between the two reduced nitrate conditions (604 DEGs, 172 overexpressed and 434 underexpressed), indicating that the cells were already acclimated to low-nitrate in the 441 µM experiment (Figure 2 E). For RCC697, only 4 genes were differentially expressed in the 220 µM condition, indicating that this nitrate reduction did not significantly affect this strain in contrast with RCC100. With only 50 µM nitrate, RCC697 exhibited an important transcriptomic response with 393 DEGs (251 underexpressed and 141 overexpressed). RCC100 and RCC697 had a similar response to the limitation of nitrate, with a total of 95 common DEGs (Supplementary Table 4).

The 6 overexpressed genes in low-nitrate environments were also overexpressed in our culture experiments when nitrate concentration was limited (Table 1). We note that the ammonium transporter was slightly differentially expressed only in the strongest nitrate depletion conditions (RCC697 50 µM). Other DEGs involved in nitrogen metabolism are described in the following paragraphs.

### Gene expression levels of *P. calceolata* RCC100 cultivated with different nitrogen compounds

*P. calceolata* RCC100 was cultivated with nitrate (882 µM, standard conditions), ammonium (882 µM), urea (441 µM) or cyanate (882 µM) for a minimum of 3 weeks with weekly transfer to fresh media. *P. calceolata* was able to grow with cyanate or urea as the sole source of nitrogen, but growth and maximal cell density were strongly limited with only ammonium as a nitrogen source (Figure 3A). To our knowledge, the uptake of cyanate in eukaryotic microalgae has only been demonstrated in *Prorocentrum*, thus growth under cyanate could involve consumption by the bacterial community and metabolite exchanges. To test this, we monitored RCC100 under cyanate in axenic versus non-axenic conditions (Figure 3B). Growth under cyanate without the bacterial community was strongly reduced, indicating that RCC100 cannot uptake cyanate for its own metabolism. Therefore, we conclude that the observed growth under cyanate relies on the bacterial community, that probably converts cyanate into another nitrogenous metabolite before assimilation by *P. calceolata*.

To determine the genes involved in the metabolic rewiring of non-axenic *P. calceolata* cultivated under different nitrogen compounds, we extracted and sequenced polyA+ RNAs in triplicate in the four conditions of Figure 3A. We obtained between 45 million and 84 million paired-end reads for each sample that were aligned on the predicted mRNAs of the *P. calceolata* RCC100 genome (Supplementary Table 3). For all conditions, triplicates were highly correlated, with Pearson correlations above 0.98 (Supplementary Figure 2B). Gene expression levels of nitrate, cyanate and ammonium conditions were more similar (Pearson > 0.95) compared to urea (Pearson between 0.73 and 0.82).

In the ammonium condition, despite strongly impacted growth, only 71 genes were underexpressed and 45 overexpressed compared to the nitrate condition. In contrast, *P. calceolata* RCC100 exhibited good acclimation to cyanate and urea and many genes were differentially expressed (921 and 308 DEGs respectively; Figure 4, A-C and Supplementary Table 5). Taken together, cyanate, urea and ammonium conditions had 65 DEGs in common, 28 of which were overexpressed and 37 underexpressed (Figure 4 D). Among the 37 underexpressed genes, 8 are involved in nitrate transport and assimilation, indicating a global down regulation of this pathway in the absence of nitrate in the medium (see discussion). We also observed down regulation of 6 LHC coding genes, suggesting a decrease in photosynthetic activity even though the alternative nitrogen source supported *P. calceolata* growth. Many overexpressed genes under alternative nitrogen sources have unknown function (13 out of 28 genes). We note the presence of 3 genes involved in protein folding (2 Heat Shock Proteins and a Cyclophilin) and 3 genes involved in protein, fatty acid and carbohydrate catabolism (a cysteine peptidase, a Phytanoyl-CoA deoxygenase and a Glycosyl hydrolase family 16), suggesting cellular stress and activation of recycling in the absence of nitrate.

**Figure 4.**
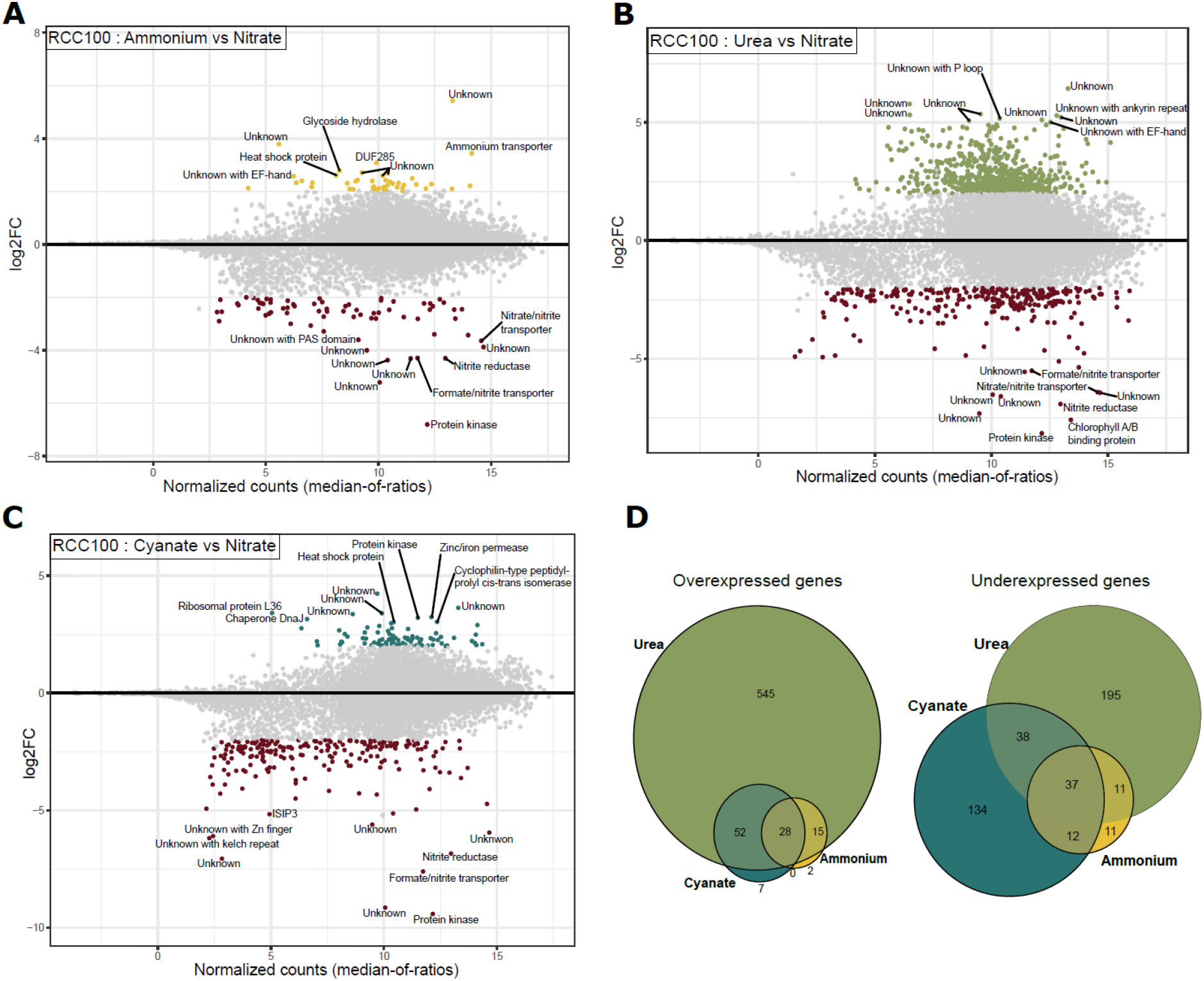
Transcriptomic response of *P. calceolata* RCC100 cultivated with different nitrogen compounds. A-C) Differentially expressed genes in 882 µM ammonium (A), 441 µM urea (B) and 882 µM cyanate (C) compared to 882 µM nitrate. Genes with p-value < 0.01 and log2FC > 2 are coloured. The function of the top 10 genes up or down regulated are indicated. D) Euler diagrams of genes overexpressed (left) or underexpressed (right) in at least one of the alternative nitrogen sources.

Only 7 genes were specifically overexpressed under cyanate (Figure 4D and Supplementary Table 5). Among these, one gene contains a domain of the transporter superfamily FepB (COG0614 ; ABC-type Fe3+-hydroxamate transport system) and carries one transmembrane domain. An enzyme carrying an alkyl-hydroperoxide reductase domain (AhpD-like, IPR029032) was also overexpressed under cyanate. This family of reductases is involved in defence against reactive oxygen species (Hong et al., 2019). The 5 other genes are a Triosephosphate isomerase (K01803-EC:5.3.1.1), a Glycosyl transferase of family 90 (PF05686), a proline iminopeptidase (K01259-EC:3.4.11.5), and genes of unknown function.

### Nitrogenous compound transport

In each family of nitrogen compound transporters, we observed genes with important variations of expression when the nitrogen source or concentration was modified (Figure 5, 6 and Supplementary Table 5). The 2 high-affinity nitrate transporters (NRT2, K02575) were downregulated when nitrate was replaced by another nitrogen source, but they were not regulated with decrease of nitrate concentration. Two formate/nitrite transporters (FNT, PF01226) were underexpressed in all alternative N sources. One FNT is a putative NAR1 transporter located on the chloroplast membrane. The other FNT was overexpressed in low-nitrate conditions *in situ* and in laboratory cultures and putatively located on lysosome/vacuole membranes, suggesting that this gene may be involved in the transportation of recycled nitrite products from intracellular vacuoles. Among the 5 ammonium transporters (Amt), one gene was overexpressed in low-nitrate conditions *in situ* and slightly overexpressed in the intermediate-NO_3_ condition, suggesting that this gene is a high-affinity transporter. This gene was also down regulated when nitrate was replaced by another nitrogen source, including ammonium. Three other Amt genes were slightly underexpressed when nitrate was depleted and might be low-affinity transporters. Interestingly, the last low-affinity transporter was strongly overexpressed in urea, cyanate and ammonium conditions, suggesting that this Amt is regulated according to the presence or absence of nitrate in the environment. One urea-proton symporter (K20989) exhibited an opposite pattern of expression in the two *P. calceolata* strains, with overexpression in low-nitrate in RCC697 and a down regulation in RCC100. The other urea transporter (PF03253) in the *P. calceolata* genome was not differentially regulated in any of our tested conditions. Finally, the uric-acid/xanthine permease (K23887) overexpressed in low-nitrate environments *in situ* was also significantly overexpressed in low-nitrate cultures and underexpressed when nitrate was replaced by cyanate or urea. This expression pattern suggests that this transporter recycles purine degradation products under nitrate starvation but does not transport extracellular nitrogenous molecules.

**Figure 5.**
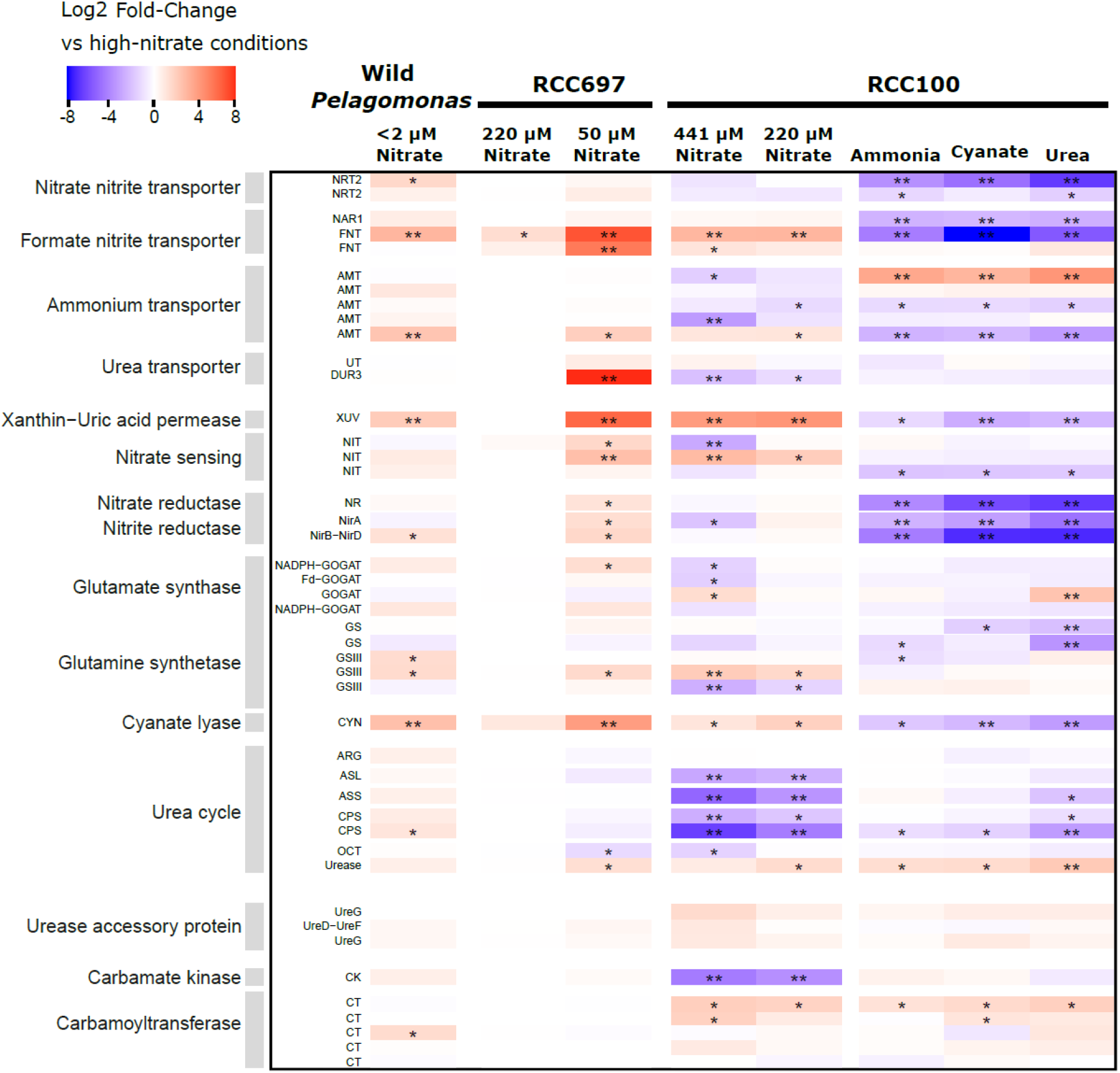
Differential expression of *P. calceolata* genes involved in nitrogen metabolism. The column named wild *Pelagomonas* represents DEGs in *Tara* Oceans samples between low-nitrate (< 2 µM) and high-nitrate (> 2 µM) environments. * = log2FC > 1 or < -1 and p-value < 0.01. **= log2FC > 2 or < -2 and p-value < 0.01.

**Figure 6.**
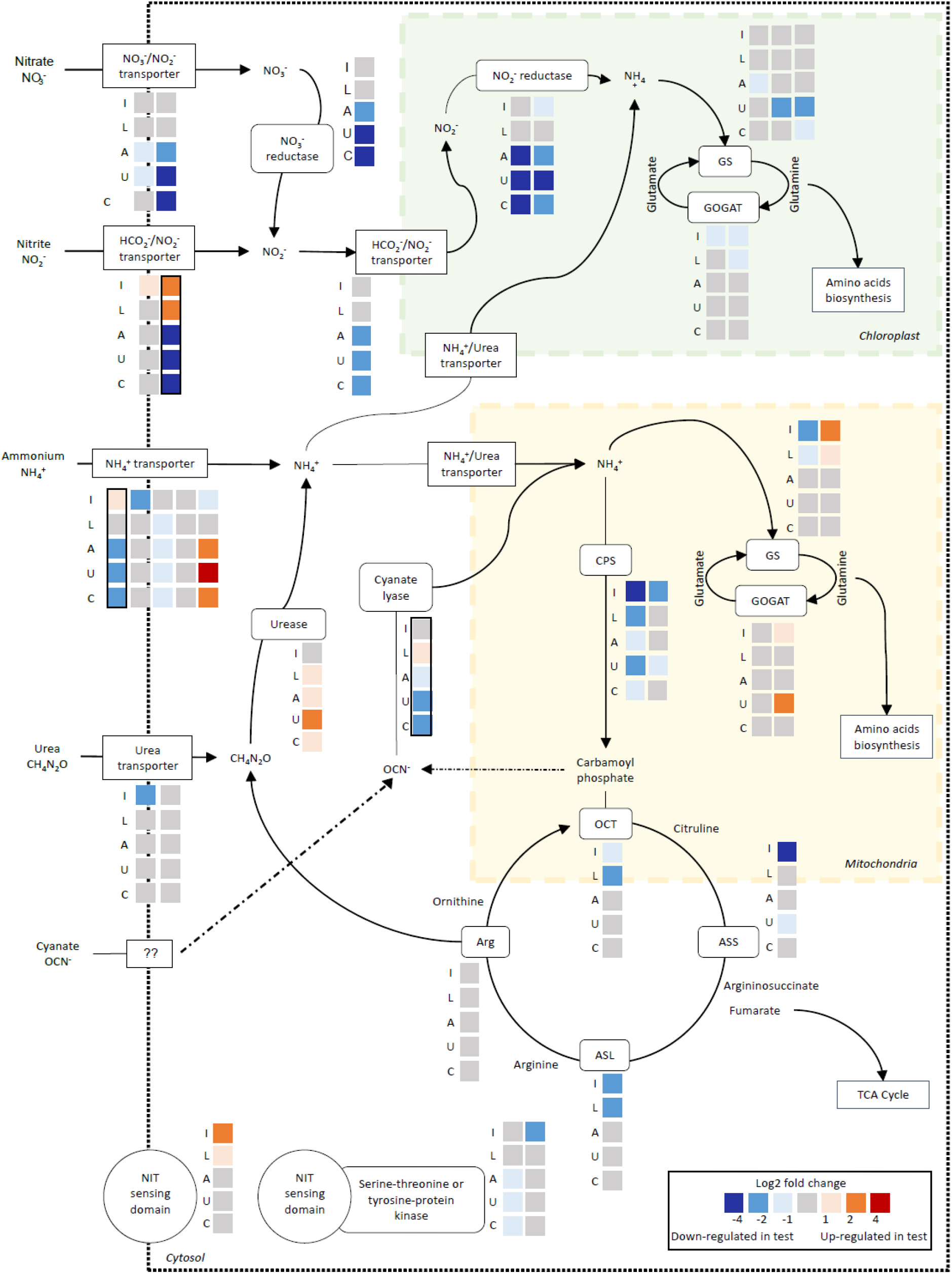
Putative nitrogen metabolism in *P. calceolata* RCC100 under low-nitrate conditions or alternative nitrogen sources. Each square represents the differential expression of one gene in one condition compared to 882 µM nitrate. Each column is one gene and rows are the different culture conditions. I: intermediate nitrate (441 µM); L: low-nitrate (220 µM); A: ammonium; U: urea; C: cyanate. GS: glutamine synthetase; GOGAT: glutamate synthase; CPS: carbamoyl-phosphate synthetase; OCT: ornithine-carbamoyl transferase; ASS: arginosuccinate synthase; ASL: arginosuccinate lyase; Arg: arginase. The colour for each square indicates if the gene is overexpressed (red) or underexpressed (blue). Genes with black lines are overexpressed in environmental low-nitrate samples.

### Nitrate sensing

Three genes carrying a nitrate/nitrite sensing domain (NIT) are present in the *P. calceolata* genome. These genes had distinct expression patterns according to nitrate concentration or source (Figure 5 and 6). The NIT-sensing gene carrying a transmembrane domain was overexpressed in low-nitrate samples, but not differentially expressed when nitrate was replaced by another nitrogen source. This gene has been shown to be overexpressed by *P. calceolata* in low-nitrate environments (Dupont et al., 2015; Guérin et al., 2022), but was not significant in our environmental DESeq2 analysis. The two other nitrate-sensing genes carry a serine–threonine/tyrosine kinase domain and might play a role in phosphorylation-based signal transduction. One of these two genes was slightly underexpressed in all alternative nitrogen sources, while the second exhibited an opposite pattern of expression in the two *P. calceolata* strains in reduced-nitrate conditions. This observation suggests that each gene has a specific role to respond to intracellular or extracellular nitrate concentration.

### Nitrate reduction and storage

The NADH-dependant nitrate reductase (NR, K10534), the NADH-dependant nitrite reductase (NirB-NirD, PF01077) and the ferredoxin-dependant nitrite reductase (NirA, K00366) had the same expression pattern with higher gene expression levels when the nitrogen source was nitrate and strong down regulation under urea, cyanate and ammonium (Figure 5 and 6). The down regulation of the nitrate reduction pathway in the absence of nitrate is coherent with previous results on many algae, including the diatom *P. tricornutum* and the pelagophyte *A. anophagefferens* (Dong et al., 2014; Smith et al., 2019). Four of the 5 genes coding for glutamine synthetase (GS) in *P. calceolata* were differentially expressed in at least one experiment. Two GS were underexpressed when the nitrogen source was urea, suggesting that urea uptake and metabolism do not require these genes. Two putatively mitochondrial GS were differentially expressed according to nitrate concentration in the laboratory experiments with opposite patterns. Four genes encode *P. calceolata* glutamate synthases (GOGAT). The putative Fd-GOGAT and one NADPH-GOGAT (GLT1, K00266), predicted to be located in the chloroplast, were slightly underexpressed in low-nitrate samples. Conversely, the putatively mitochondrial GOGAT was slightly overexpressed in low-nitrate experiments (Figure 5 and 6).

### Cyanate lyase, urease and urea cycle

As in low-nitrate environmental samples, the *P. calceolata* cyanate lyase gene was slightly overexpressed in our low-nitrate experiment in both strains (Table 1). Surprisingly, under alternative nitrogen sources, including cyanate, the cyanate lyase was underexpressed. This result shows that cyanate lyase is not involved in assimilation of extracellular cyanate and could instead be involved in nitrogen recycling from intracellular molecules (see discussion). *P. calceolata* encodes one urease (URE, K01427) and three urease accessory proteins: one UreD-UreF (K03190;K03188) and two UreG (PF02492). The urease was slightly overexpressed in low-nitrate samples and overexpressed in alternative nitrogen sources, especially urea. This result indicates that extracellular urea as well as intracellularly produced urea are hydrolysed into ammonia by the urease. The urea cycle in *P. calceolata* is composed of two carbamoyl phosphate synthetases (CPS), 1 ornithine carbamoyltransferase (OCT), 1 arginosuccinate synthase (ASS), 1 arginosuccinate lyase (ASL) and 1 arginase (Figure 4). Except for the arginase, all genes involved in the urea cycle were overexpressed in high-nitrate conditions. Together with strong overexpression of the putatively mitochondrial GOGAT, this pattern indicates removal of excess ammonia from the cell through the urea cycle.

## Discussion

### *P. calceolata* has common patterns of low-nitrate acclimation in situ and in culture experiments

In this study we analysed the nitrogen metabolism of *Pelagomonas in situ* with *Tara* Oceans datasets and in laboratory culture experiments on two strains of *P. calceolata* under different nutrient conditions. Although we only identified 6 genes significantly differentially expressed in low-nitrate environments *in situ*, 5 of them were also significantly differentially expressed in low-nitrate conditions in the laboratory. This result is remarkable for two reasons. Firstly, the strains used in this study have been cultivated for several decades in nitrate-rich conditions without environmental stress. Although extended periods of cultivation may expose strains to genomic modifications (genome rearrangements, reduction, selection), the regulation of several genes in response to low-nitrate has evidently been conserved compared to wild *Pelagomonas*. Secondly, the minimal amount of nitrate used in the laboratory to obtain sufficient *P. calceolata* growth(50 µM) was one hundred times greater than the concentration measured in the oceans where *P. calceolata* is relatively abundant (below 0.5 µM). Despite this important difference, the transcriptomic pattern in response to nitrate starvation was similar.

Because of the relatively low abundance of cells in the environment, metatranscriptomes generally provide access only to highly expressed genes. Therefore, laboratory experiments are needed to observe fine-scale variations in genes with low expression levels, providing a more detailed picture of metabolic processes. In addition, given the numerous environmental parameters affecting gene expression levels in the environment, it is often difficult to dissociate the effects of different parameters in order to assess specific responses. In our analysis, we used the large diversity of environments sampled during the *Tara* cruise to get among low nitrate samples a large range of all other parameters (temperature, salinity, iron concentrations,…). In this manner, the risk of environmental variables correlated with nitrate concentration is limited and the differential expression analysis will only identify genes that were directly affected by nitrate concentrations. In addition, in environmental metatranscriptomes generated across large geographical areas it can be challenging to determine whether gene expression variations are the result of acclimation (reversible short-term transcriptomic regulation) or adaptation (long-term selection). Complementing *in situ* analysis with culture-based transcriptomics allowed us to conclude that for genes that are differentially expressed in both situations, variations in *in situ* expression levels are likely the result of short-term acclimation.

### *P. calceolata* is genetically adapted to consume organic nitrogen compounds

We observed growth of *P. calceolata* under nitrate, urea or cyanate as the sole nitrogen source, demonstrating that this species is adapted to consume organic nitrogen compounds, like many other microalgae. The related pelagophyte *A. anophagefferens*, for example, exhibits enhanced growth when cultivated with urea compared to other nitrogen sources such as nitrate or ammonium (Berg et al., 1997). The transcriptomic profile of *P. calceolata* was significantly modified when cultured with organic nitrogen compounds, but stress response genes were not overexpressed. Our results allow us to distinguish the common transcriptomic response in the presence of urea, cyanate or ammonium from the specific responses to each nitrogen source. The main common response in the absence of nitrate was down-regulation of elements of the nitrate assimilation pathway, such as formate-nitrite and nitrate-nitrite transporters, as well as nitrate and nitrite reductases. This pathway was not under-expressed when nitrate concentration was reduced, suggesting that residual nitrate in our low-nitrate conditions was sufficient to maintain the pathway. The presence of urea, which is the best studied alternative nitrogen source, negatively affects plastidic nitrogen assimilation in the pelagophyte strain CCMP2097 (Terrado et al., 2015). In *A. anophagefferens*, the use of urea as a nitrogen source was observed to trigger up-regulation of genes involved in protein, amino acid, spermine and sterol synthesis (Dong et al., 2014), but these functions were not up-regulated in our experiments with *P. calceolata*.

### Recycling of intracellular nitrogenous compounds is the dominant response under low-nitrate conditions

This study shows that under low-nitrate, the main strategy is the recycling of intracellular nitrogenous compounds such as amino acids and nucleotides. A gene coding a xanthine/uracil/vitamin C permease (XUV), involved in the transport of purine degradation products, is overexpressed both in environmental and experimental low-nitrate conditions. Overexpression of XUV under low-nitrate conditions has previously been observed in several microalgae, including the pelagophyte *A. anophagefferens* and the haptophyte *Prymnesium parvum* (Dong et al., 2014; Liu et al., 2015; Wurch et al., 2014). The expression of this permease in *P. calceolata* seems to also be linked to catabolism of purines as a recycling mechanism in low-N environments and does not reflect the presence of purines in the environment. Enzymes necessary for the conversion of xanthine into urea and ammonia are present in the *P. calceolata* genome, but were not over-expressed in low-N conditions. This pathway is unlikely to be involved in purine recycling in *P. calceolata*, in contrast to *A. anophagefferens* when supplied with xanthine (Gann et al., 2022). Recent research highlights that many microalgae including, ochrophytes, are able to store nitrogen in purine crystals (Pilátová et al., 2022). Over-expression of purine permease in *P. calceolata* under low-nitrate could be a sign of nitrogen reallocation from crystals, but crystalline inclusions have not been reported in pelagophytes to date.

### Role of cyanate lyase in *P. calceolata*

Among eukaryotic microalgae, growth with cyanate as the sole nitrogen source has only previously been shown in the dinoflagellate *Prorocentrum donghaiense*, at the expense of a lower growth rate (Hu et al., 2012). It has been suggested that phytoplankton overexpressing cyanate lyase in low-nitrate environments are therefore capable of cyanate uptake and metabolism (Dong et al., 2014; Mao et al., 2022). Although, *P. calceolata* cyanate lyase is overexpressed in low-nitrate environments (*in situ* and in culture), we have shown that the bacterial community is required for *P. calceolata* to thrive under cyanate and that the cyanate lyase gene is under-expressed in this case. In consequence, cyanate lyase seems to not be involved in external cyanate metabolism and should not be used as a marker of cyanate uptake. In agreement with our results, cyanate supports the growth of several ascomycete species despite the absence of cyanate lyase coding genes in their genome (Linder, 2018). Conversely, several yeasts with cyanate lyase genes were unable to grow under cyanate, supporting the hypothesis that this gene is not involved in the assimilation of external cyanate in eukaryotes in contrast to prokaryotes. In *P. calceolata*, like all other eukaryotes, the bacterial cyanate transporter (CynX) is not conserved (Widner & Mulholland, 2017). Among the 7 genes specifically over-expressed in the sole presence of cyanate, a single gene carries homologies with a protein transporter domain (ABC-type Fe3+-hydroxamate). Since this gene has only one transmembrane domain, it is probably not involved in cyanate uptake.

The function of cyanate lyase in *P. calceolata* is likely detoxification of intracellularly produced cyanate. Cyanate can be generated through rapid decomposition of carbamoyl-phosphate (Anderson et al., 1990; Ter-Ovanessian et al., 2021) or slow decomposition of urea (Dirnhuber & Schütz, 1948). *P. calceolata* recycles amino acids and proteins when nitrate supply is limited. Many enzymes are involved in the catabolism of metabolites in this process, including carbamoyltransferases (K00612) that can produce carbamoyl-phosphate (CP). Two of the five genes coding for carbamoyltransferases were over-expressed in low-nitrate conditions in RCC100 (Figure 5). We hypothesise that these enzymes increase intracellular concentration of CP, which is in principle taken up by the urea cycle. However, under low-nitrate conditions the urea cycle is down-regulated, which could lead to an increase of intracellular cyanate, requiring cyanate lyase to detoxify the cell and produce ammonia. In complement to the work of Sato, Hashihama, and Takeda 2023 and the phylogeny of Mao et al. 2022, our results suggest that the removal of intracellular cyanate rather than external cyanate metabolism as an alternative N source is the main role of cyanate lyase in eukaryotic microalgae.

## Data Accessibility

The transcriptomic data generated in this study have been deposited in the European Nucleotide Archive database under accession code PRJEB74085. Gene expression levels of *P. calceolata* are on Zenodo for laboratory experiments (https://zenodo.org/uploads/12582059) and Tara Oceans metatranscriptomes (https://doi.org/10.5281/zenodo.6983364). All codes for the bioinformatic workflow are provided at: https://github.com/institut-de-genomique/PelagomonasNitrogenMetabolism.

## Author Contributions

N.G and Q.C conceived and planned this study with the support of P.W. A.T set up the culture chamber with important advice from P.G and I.P. N.G and C.S performed *P. calceolata* cultures with the strong support of C.O and L.B. B.V, E.B and G.M did RNA extractions, library preparations and sequencing coordinated by K.L. N.G and C.S carried out all bioinformatics analysis supervised by Q.C. N.G and Q.C wrote the paper. All authors contributed to the manuscript preparation and approved the final version of the paper.

## Acknowledgments

We thank the commitment of the following people who made this work possible: the Genoscope/CEA, Paris-Saclay University and the CNRS; members of the Roscoff Culture Collection for maintaining *Pelagomonas* strains; Claude Scarpelli for support in high-performance computing at Genoscope. We acknowledge the financial support of the ANR (ANR-22-CE20-0012; ANR20-CE02-0025) and FRANCE GENOMIQUE (ANR-10-INBS-09–08). We also thank the *Tara* Expedition Foundation and their partners for the organization of marine scientific expeditions (http://oceans.taraexpeditions.org).

