## Supplementary Figure for "Nitrogen metabolism in the picoalga *Pelagomonas calceolata*: disentangling cyanate lyase function under different nutrient conditions"

### Supplementary information

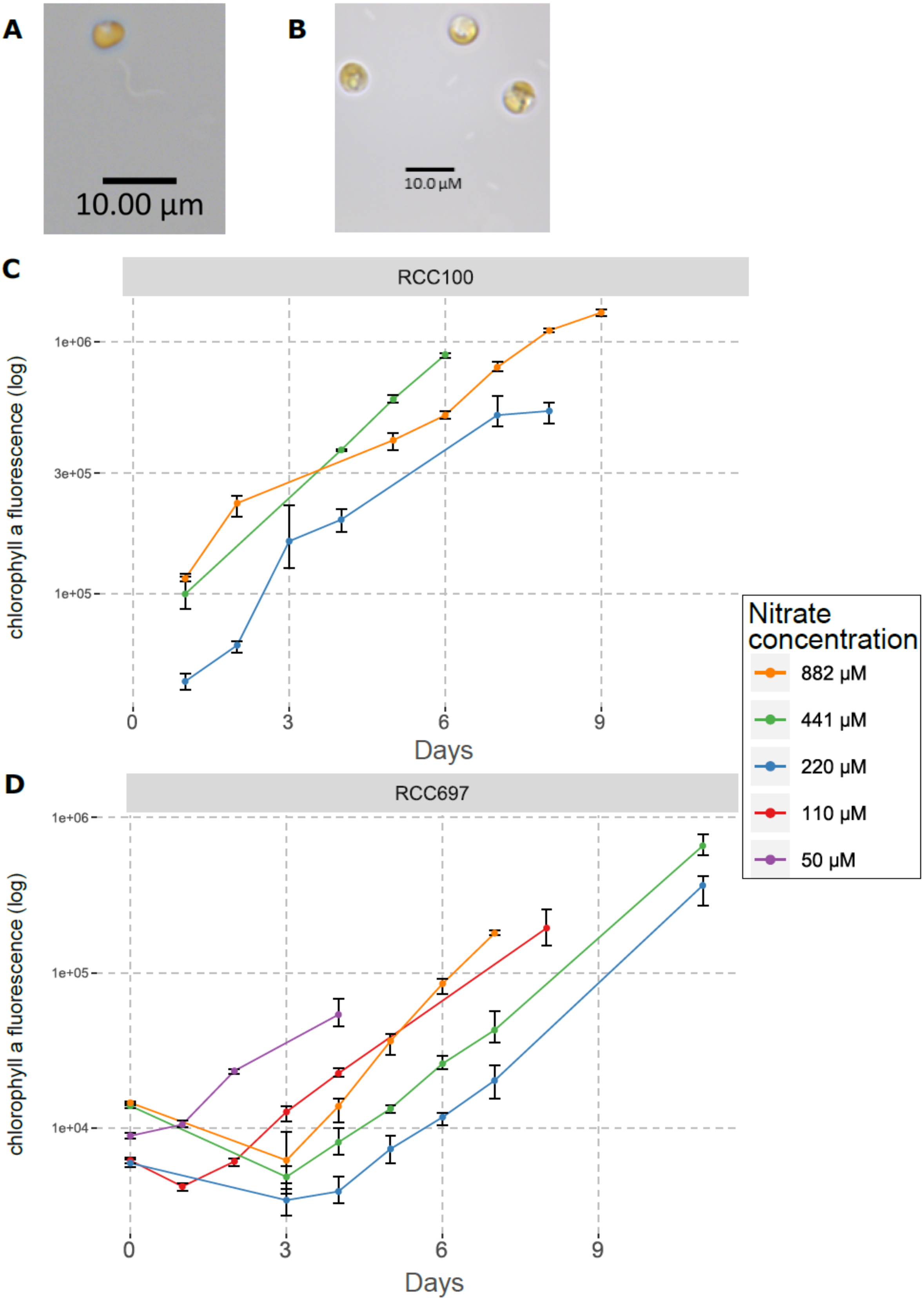


**Supplementary Figure 1:** A) *P. calceolata* RCC100, light microscopy. B) *P. calceolata* RCC697, light microscopy. C and D) *P. calceolata* RCC100 (C) and RCC697 (D) growth with different nitrate concentrations. The chlorophyll a fluorescence is used as a proxy for cell concentration (arbitrary unit). *P. calceolata* cells were harvested for RNA extraction at the last time point of each condition.

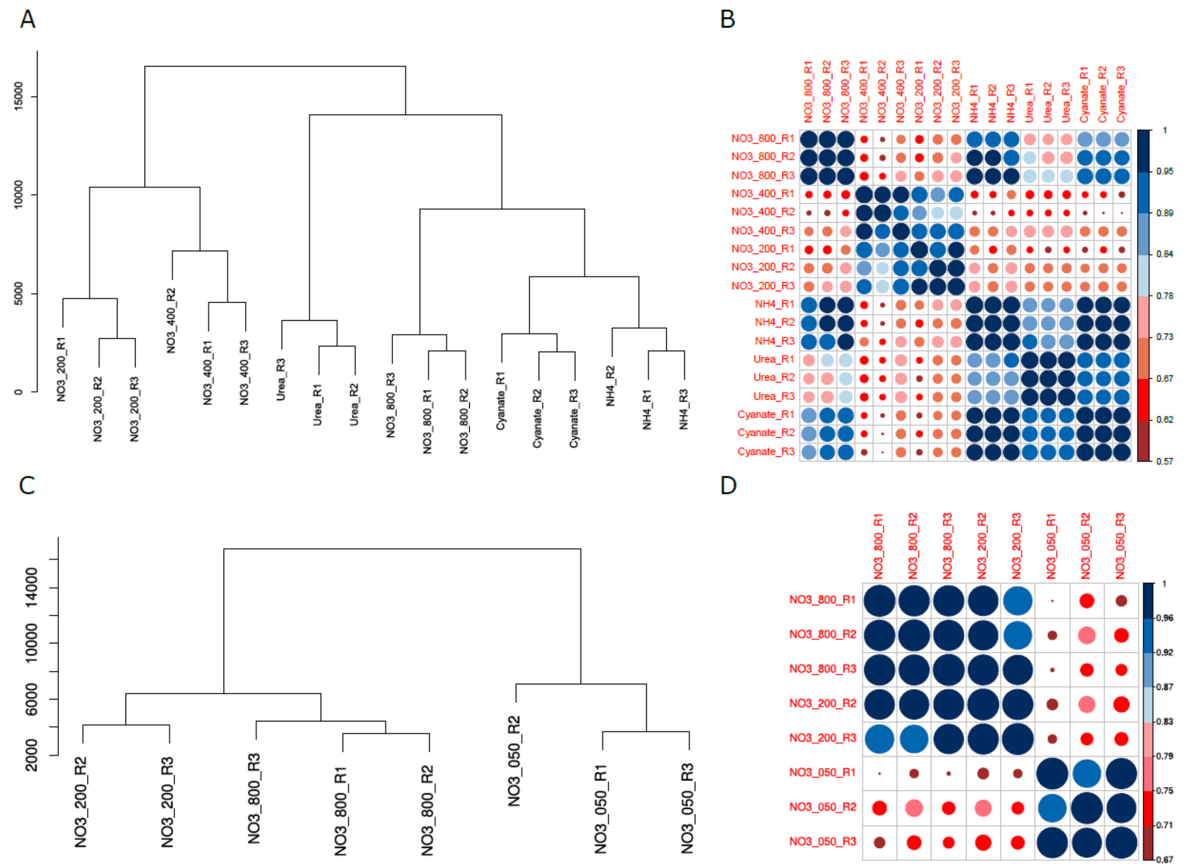


**Supplementary Figure 2: Gene expression levels of *P. calceolata* RCC100 and RCC697 under different nitrate concentrations.** A and C) Hierarchical clustering of the gene expression levels of *P. calceolata* RCC100 (A) and RCC697 (C). B and D) Pearson's r correlations of gene expression levels between each pair of samples for *P. calceolata* RCC100 (B), and RCC697 (D). Each replicate is processed independently and is indicated by R1 to R3 for each condition
